# Lhcb9-dependent photosystem I structure in moss reveals evolutionary adaptation to changing light conditions during aquatic-terrestrial transition

**DOI:** 10.1101/2023.01.18.524345

**Authors:** Haiyu Sun, Hui Shang, Xiaowei Pan, Mei Li

## Abstract

In plants and green algae, light-harvesting complexes I and II (LHCI and LHCII) constitute the antennae of photosystem I (PSI), thus effectively increasing the cross-section of the PSI core. The moss *Physcomitrium patens* (*P. patens*) represents a well-studied evolutionary intermediate between green algae and flowering plants. *P. patens* possesses at least three types of PSI with different antenna sizes. The largest PSI form (*Pp*PSI-L) exhibits a unique organization found neither in flowering plants nor in algae. Its formation is mediated by the *P. patens*-specific LHC protein, Lhcb9. While previous studies revealed the overall architecture of the *Pp*PSI-L, its assembly details and the relationship between different *Pp*PSI types remain unclear. Here, we report a high-resolution structure of the *Pp*PSI-L. We identified 14 PSI core subunits, one Lhcb9, one phosphorylated LHCII trimer, and eight LHCI monomers arranged as two belts. Our structural analysis established the essential role of Lhcb9 and the phosphorylated LHCII in stabilizing the complex. In addition, our results suggest that *Pp*PSI switches between three different types, which share identical modules. This feature may contribute to the quick and dynamic adjustment of the light-harvesting capability of PSI under different light conditions.

## Introduction

Oxyphototrophs (plants, algae and cyanobacteria) utilize light energy to convert carbon dioxide and water into organic compounds, in the process of which oxygen is released[1, 2]. All of these products are indispensable for most life forms on Earth. As oxyphototrophs, plants are responsible for approximately half of the primary production through photosynthesis, and they emerged approximately 450 million years ago, when green lineage (green algae and plants) transited from an aqueous to a terrestrial environment[3]. Bryophytes, which comprise liverworts, mosses and hornworts, distinguished themselves in this important evolutionary event as the first types of the terrestrial green plants, thus represent evolutionary intermediate between aquatic green algae and terrestrial flowering plants[4, 5]. One moss, *Physcomitrium patens* (*P. patens*, also known as *Physcomitrella patens*), has been used extensively as a model plant in a wide range of research fields since its genome was completely sequenced [5, 6].

Oxygenic phototrophs possess two photosynthetic machineries known as photosystems I and II (PSI and PSII), which are large assemblies localized within the thylakoid membranes[7]. Both photosystems consist of a core complex responsible for light-induced charge separation and electron transfer processes, and a peripheral antenna system serving to increase the absorption cross-section of the core. The antenna system is composed of various light-harvesting complexes (LHCs), which are encoded by a large gene family called *lhca* and *lhcb* genes[8]. The LHCIs (Lhcas, *lhca* gene products) are mostly present as monomers and associated with the PSI core, whereas the LHCIIs (Lhcbs, *lhcb* gene products) primarily bind to the PSII core and are classified into two types, namely the major LHCII trimers and the minor LHCII monomers[8-10].

While green algae and plants contain a highly conserved core complex, they differ in the size and organization pattern of peripheral antenna system for both photosystems, presumably to better adapt to different light conditions in their respective environments [11]. For example, in a unicellular green alga *Chlamydomonas reinhardtii* (*C. reinhardtii*), nine distinct Lhca (Lhca1-9) proteins have been identified[12, 13]. The core has been shown to bind up to ten of these Lhcas, thus forming a large PSI-LHCI complex[14, 15]. In comparison to green algae, plants possess fewer Lhca proteins and the PSI complex with smaller antenna system[16, 17], which might be a result of acclimation to the higher light intensity at terrestrial environments. The model plant *Arabidopsis thaliana* (*A. thaliana*) contains six *lhca* genes (*lhca*1-6); however, *lhca*5 and *lhca*6 are expressed at the sub-stoichiometric level[18]. Four Lhca proteins are arranged in the order of Lhca1-a4-a2-a3, and attach to the core as one LHCI belt in plant PSI [19]. Lhca5 is able to occupy the Lhca4 position in the Lhca4-deficient mutant, forming a PSI-LHCI similar to the canonical one albeit with lower amount of the complex[20-22].

The moss *P. patens* only contains *lhca*1-3 and *lhca*5 genes, while *lhca*5 shows much lower expression level than *lhca*1-3[23-25]. Furthermore, *P. patens* contains a unique LHC protein, namely Lhcb9, which is similar to Lhcb proteins in sequence, but possesses the red chlorophyll (Chl) characterized for Lhca proteins[23, 25, 26]. Lhcb9 has two isoforms (Lhcb9.1 and Lhcb9.2) with a sequence identity of approx. 80%[23, 27, 28]. In addition to Lhcb9, most of *P. patens* PSI subunits contain more than one isoform, including Lhca1-3[23, 25]. The high number of isoforms in *P. patens* is likely due to a whole-genome duplication[5, 23, 25, 29]. An earlier report showed that several PSI subunits show altered expression profile in different development stages (protonemata and gametophore) of *P. patens*, suggesting their role in fine-tuning the PSI function[24].

In addition to containing multiple isoforms of PSI and LHCI subunits, *P. patens* also possesses several types of PSI complex which differ in the size and composition of the antenna system. The smallest *Pp*PSI form resembles the PSI-LHCI found in vascular plants, contains one PSI core associated with four Lhca proteins (hereafter termed *Pp*PSI-S), although *P. patens* lacks the *lhca*4 gene[24, 30, 31]. Previous structural analysis of *Pp*PSI-S suggested that Lhca5 occupies the Lhca4 position[30], whereas a recent *Pp*PSI-S structure with higher resolution revealed that two different Lhca2 paralogues (Lhca2b and Lhca2a) are located at the two middle positions of the LHCI belt, resulting in the LHCI belt with Lhca1-a2b-a2a-a3 organization[31]. Medium-sized PSI complexes, including the state transition complex PSI-LHCI-LHCII[24, 32, 33], were previously observed in *P. patens*. State transitions are a short-term acclimation mechanism that responds to the redox state of plastoquinone pool and balances the energy distribution between two photosystems in plants and green algae[34, 35]. Under state 2 conditions, a portion of LHCII trimers is phosphorylated and binds to the PSI core, forming the PSI-LHCI-LHCII complex that increases the excitation energy flowing to the PSI[36-39]. Structures of the PSI-LHCI-LHCII complex from *Zea mays* and *C. reinhardtii* (*Zm*PSI-LHCI-LHCII and *Cr*PSI-LHCI-LHCII) were previously reported[38, 40], showing that the phosphorylated LHCII is essential for the complex formation. The formation of *Pp*PSI-LHCI-LHCII complex also depends on the phosphorylated LHCII[32], however, direct structural information of *Pp*PSI-LHCI-LHCII remains absent.

The largest *Pp*PSI complex (hereafter termed *Pp*PSI-L) is mediated by the *P. patens*-specific protein Lhcb9[27, 28, 33]. In addition to the plant-type PSI-LHCI and Lhcb9, the *Pp*PSI-L complex contains one LHCII trimer and additional four LHC proteins[27, 33]. Deletion of Lhcb9 results in the dissociation of the *Pp*PSI-L complex[27, 28, 33]. Moreover, formation of this complex is greatly reduced in a *P. patens* mutant lacking LHCII kinase STN7[32]. These results demonstrated that both Lhcb9 and phosphorylated LHCII are indispensable for the *Pp*PSI-L formation. The phosphorylated LHCII associated with the PSI-LHCI complex is essential for state transitions; however, deletion of Lhcb9 showed little effect on the state transition level of *P. patens*, indicating that the *Pp*PSI-L complex is not involved in this regulatory mechanism[28, 33]. Presumably, the complex is critical for increasing the light harvesting capability of the *P. patens* PSI at low light, and disassembles under prolonged high light conditions[27, 33]. Currently available low-resolution electron microscopy (EM) maps of *Pp*PSI-L have established critical aspects of the architecture of *Pp*PSI-L and the potential binding position of Lhcb9[27, 33]. However, the exact composition of LHC proteins, the detailed inter-subunit interactions and the potential excitation energy transfer (EET) pathways of the *Pp*PSI-L complex remain unclear. Here, we report the 2.87 Å resolution cryo-EM structure of *Pp*PSI-L, providing information about the protein composition and assembly, as well as the sophisticated network of chlorophylls mediating the efficient EET within this complex.

## Results

### Overall structure of *Pp*PSI-L complex and the *Pp*PSI-S moiety

We purified both *Pp*PSI-L and *Pp*PSI-S complexes from wild-type moss *P. patens* (Fig. S1) grown under low light conditions in the presence of glucose, according to a method described in a previous report[28, 33]. Compared to *Pp*PSI-S, *Pp*PSI-L shows higher absorption at the regions characterized for Chl b molecules (Fig. S1c), indicating that it binds Lhcb proteins. Immunoblotting confirmed that the *Pp*PSI-L complex contains Lhcb9 and phosphorylated LHCII, both of which are absent in our *Pp*PSI-S samples (Fig. S1e). To identify whether *Pp*Lhca5 is present in our *Pp*PSI-S and *Pp*PSI-L complexes, we performed mass spectrometry (MS) analysis on each protein band separated by SDS-PAGE (Fig. S1b), and found that Lhca5 is absent in both samples (Source data). We then solved the cryo-EM structure of *Pp*PSI-L complex at an overall resolution of 2.87 Å (Fig. S2, Table S1). This structure showed that *Pp*PSI-L complex contains a PSI core, one Lhcb9, one LHCII trimer, and eight LHCI monomers arranged as two LHCI belts (hereafter termed as the inner and outer belt), each composed of four Lhca proteins (Figs. 1, 2a). The inner belt associates with the PSI core at the PsaG-F-J-K side, resembling the *Pp*PSI-S and PSI-LHCI complex from vascular plant [30, 31, 41]. The LHCII trimer attaches to the PSI at the PsaO side, similar to that of *Zm*PSI-LHCI-LHCII complex. The outer belt associates with the inner belt and LHCII through two terminal Lhca proteins, encircling a central space where Lhcb9 is located. Interestingly, the membrane spanning region of *Pp*PSI-L complex is curved. The LHCII trimer and the outer LHCI belt display opposite curvatures, and are pivoting towards the stromal and luminal side, respectively, in relation to the core (Fig. 1b). The curved arrangement of the complex may be related to the thylakoid membrane architecture of *P. patens*[42].

**Fig. 1.**
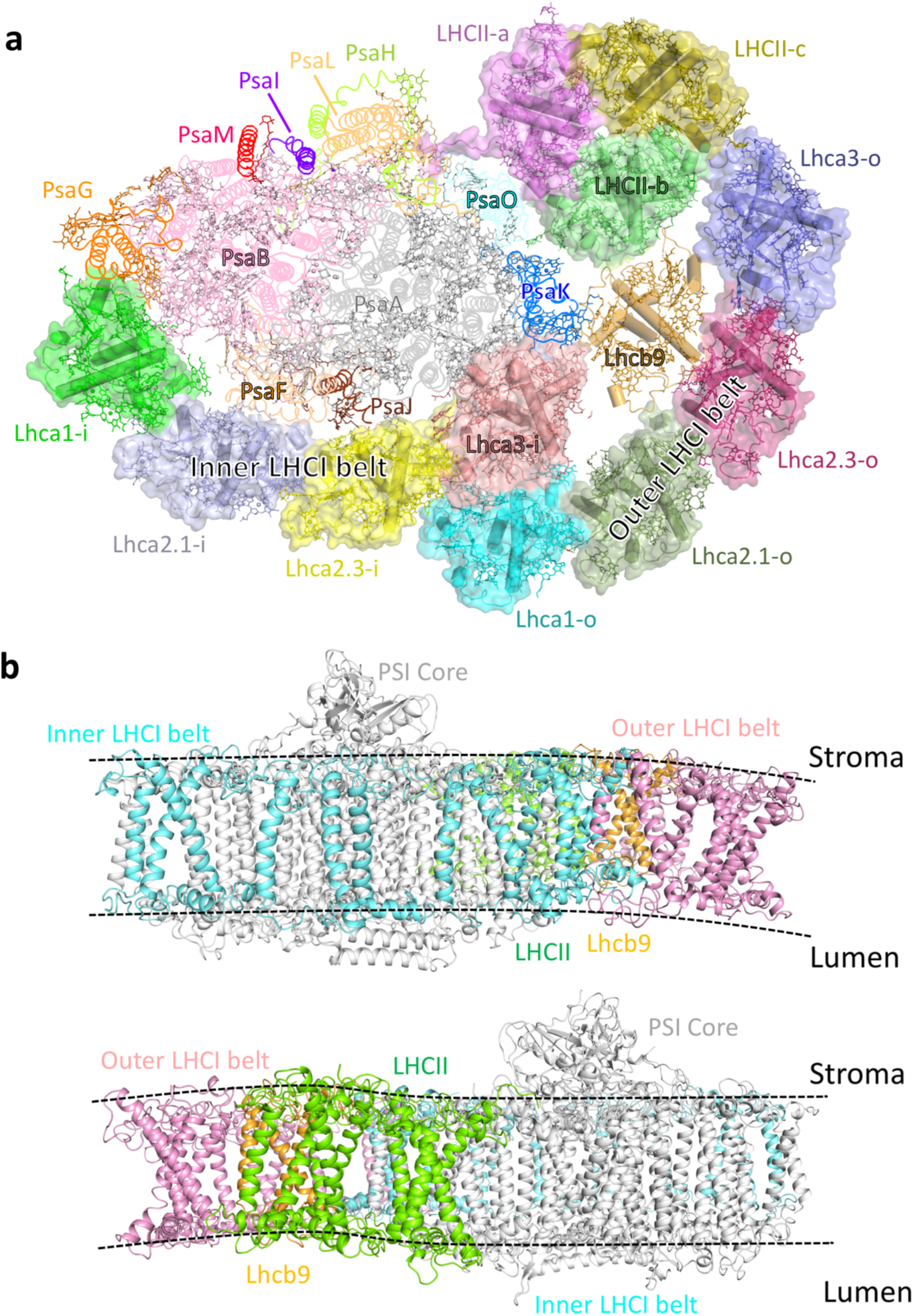
Overall structure of *Pp*PSI–L complex. **a**, Stromal side view of the *Pp*PSI–L complex. The core subunits and LHC proteins are shown in ribbon and cartoon mode, respectively. The LHCII trimer, the inner and outer LHCI belts are further displayed in surface. Pigments, lipids and other co-factors are shown in stick mode. Membrane-extrinsic subunits PsaC, PsaD and PsaE are omitted for clarity, while other subunits are shown in different colors and labelled. **b**, Side view of the *Pp*PSI–L complex. PSI core, inner LHCI belt, outer LHCI belt, LHCII, Lhcb9 are shown in different colors and labelled. Black dashed lines indicate the curved membrane spanning region of *Pp*PSI-L complex.

**Fig. 2.**
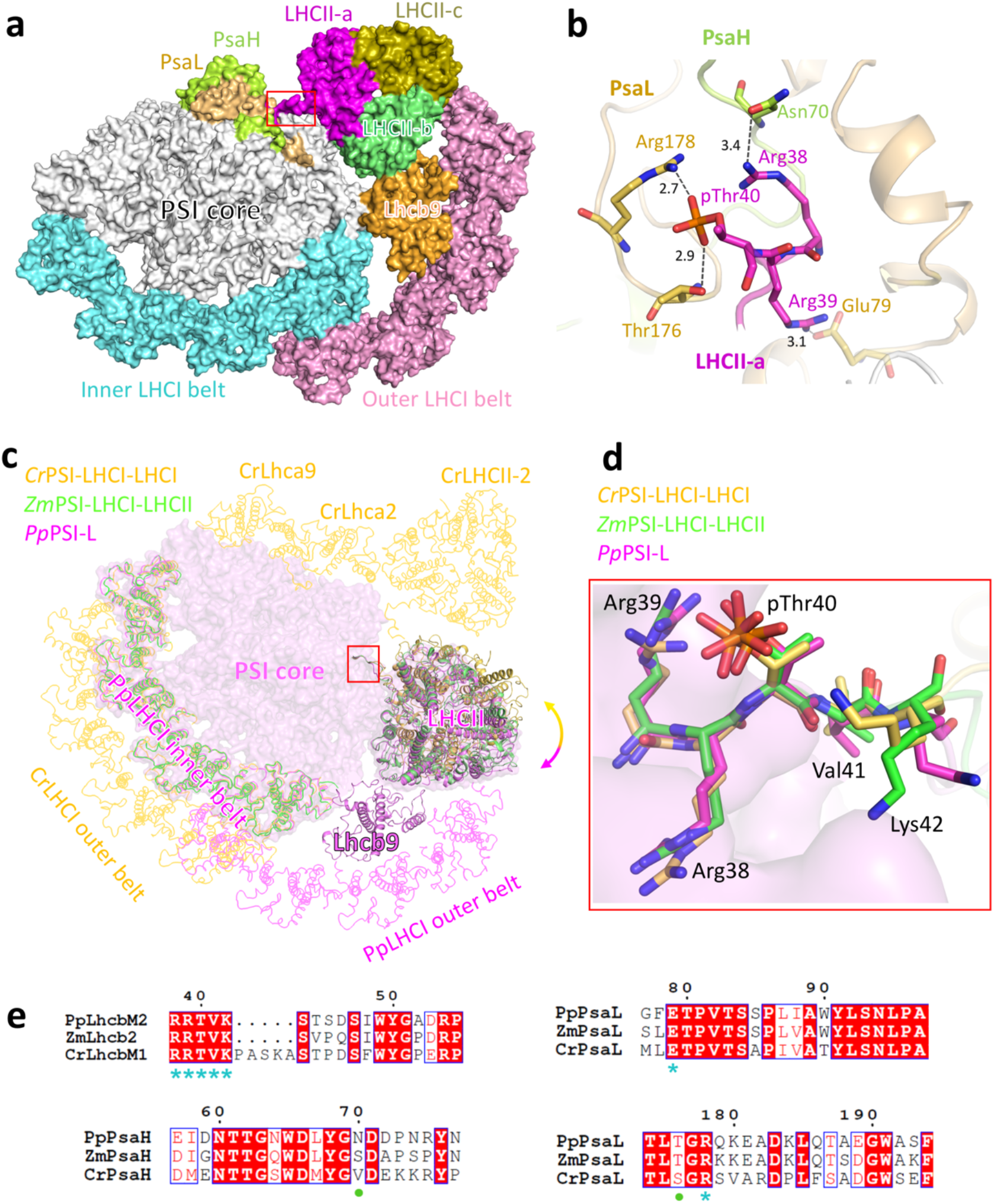
Interactions between the LHCII trimer and the PSI core. **a**, Surface representation of the *Pp*PSI-L complex. The phosphorylated N-terminal region of LHCII-a (LhcbM2) is highlighted by a red box. **b**, Detailed interactions between the N-terminal region of LHCII-a and PSI core subunits PsaL and PsaH. The hydrogen bond interactions are represented with dashed lines and the distances (Å) are labelled. **c**, Structural superposition of *Pp*PSI-L (magenta) with *Cr*PSI-LHCI-LHCI (yellow, PDB code 7DZ7) and Z*m*PSI-LHCI-LHCII (green, PDB code 5ZJI) viewed from the stromal side. Colored arrows indicate the rotational direction of LHCII in *Pp*PSI-L and LHCII-1 in *Cr*PSI-LHCI-LHCII with respect to the corresponding LHCII in *Zm*PSI-LHCI-LHCII. The red box highlights the N-terminal regions of LHCII monomers in the three structures. The PSI core, inner LHCIs and LHCII in *Pp*PSI-L are also shown in surface to represent the *Pp*PSI-LHCI-LHCII complex. **d**, Structural comparison of N-terminal five residues (shown as sticks) of *Pp*LhcbM2 (carbon atoms colored magenta), *Cr*LhcbM1 (carbon atoms colored yellow) and *Zm*Lhcb2 (carbon atoms colored green). The PSI core is shown in surface mode. **e**, Sequence alignment of the N-terminal regions of *Pp*LhcbM2, *Zm*Lhcb2 and *Cr*LhcbM1, as well as the interaction-related regions of PsaL and PsaH subunits from three species. Conserved residues involved in inter-subunit interactions are indicated by blue asterisks. Green dots indicate residues in *Pp*PSI-L involved in LHCII interaction but not completely conserved during evolution.

Our *Pp*PSI-L structure contains 26 protein subunits and 366 cofactors (Fig.1, Table S2). The PSI core possesses 14 subunits, including PsaM, but absent of PsaN, consistent with previous *P. patens* genome analysis[5, 23, 24]. We identified the four Lhca proteins in the inner belt based on our cryo-EM map. Lhca1 and Lhca3 are located at two ends of the inner belt, close to PsaG and PsaK, respectively. Two Lhca2 isoforms, Lhca2.1 and Lhca2.3 (corresponding to Lhca2b and Lhca2a in 7KSQ) [31], occupy the position of Lhca4 and Lhca2 in plant PSI-LHCI (Fig. 1a, Fig. S3). The assignment of Lhca2.1 instead of Lhca5 in the second position of the LHCI belt is in agreement with our MS result. The *Pp*PSI-S moiety in our *Pp*PSI-L complex is almost identical to the recently reported *Pp*PSI-S structure (7KSQ), with the same Lhca composition and arrangement (Fig. S4). These results suggested that the *Pp*PSI-L complex is assembled by the *Pp*PSI-S bound with additional LHC proteins, but not a newly assembled PSI core with various LHCs. This assembly pattern may be beneficial for rapid regulation of the light harvesting capacity of *P. patens* PSI simply by controlling the association/dissociation of the additional LHCs with/from the *Pp*PSI-S.

### LHCII binds PSI through its phosphorylated N-terminal region

Our structural analysis showed that the *P. patens* LHCII trimer exhibits the same structural features as those from plants and green algae, possesses three transmembrane helices (B, C, A from N-to C-terminus) and two short helices (E, D) at the luminal side (Fig. S5a). Each LHCII monomer binds 14 chlorophylls (Chl 601-614, nomenclature according to [43]) and four carotenoids located at the conserved L1, L2, N1 and V1 site (Fig. S5a). In the *Pp*PSI-L structure, one LHCII monomer is clearly phosphorylated (Fig. S5b,c) at its N-terminal tail, which interacts with the PSI core (Fig. 2a). The structural observation that LHCII is phosphorylated is in agreement with the immunoblotting results from both our experiment (Fig. S1e) and previously reported data [32, 42]. We were unable to identify the specific isoform of the trimer solely based on the density map, since *P. patens* contains 14 LhcbM isoforms, all of which are involved in the trimer formation, and which display highly similar sequences (identity higher than 90%). However, by analyzing the feature of residues Tyr^57^ instead of Phe^57^ of our cryo-EM map, we found the candidates for the phosphorylated monomer to be LhcbM1, LhcbM2, LhcbM4, LhcbM5 and LhcbM13 (Fig. S5d). Moreover, previous analysis of LhcbMs phosphorylation revealed that LhcbM2, LhcbM4/8 and LhcbM13 are phosphorylated at their N-terminal region in large *Pp*PSI complexes[32]. Furthermore, our MS analysis of the *Pp*PSI-L sample detected unique peptides belonging to LhcbM1 and LhcbM2. The two LhcbM isoforms were also identified in previous studies of *Pp*PSI-L complex [27, 33]. Based on these results, we propose that LhcbM2 constitutes the monomer that is phosphorylated and directly interacts with the PSI. We modeled the three monomers using LhcbM2 sequence and termed them LHCII-a (phosphorylated LhcbM2), LHCII-b (the monomer adjacent to Lhcb9) and LHCII-c (the outside monomer) (Fig. 1).

In addition to the hydrophobic interactions at the membrane region, LHCII associates with the PSI mainly through the first three residues of LhcbM2 (^38^RRt^40^) at the stromal side (Fig. 2b). The pThr^40^ of LhcbM2 forms an ionic bond with Arg^178^ and is hydrogen bonded with Thr^176^ of PsaL, the first two Arg residues interact with residues from PsaL and PsaH. These interactions are critical for stabilizing the association of LHCII with the PSI core. When we superposed our *Pp*PSI-L structure with *Zm*PSI-LHCI-LHCII and *Cr*PSI-LHCI-LHCII structures, we found that the LHCII trimer in the three structures is located at a similar position. In addition, their phosphorylated LHCII monomer (*Pp*LhcbM2, *Zm*Lhcb2 and *Cr*LhcbM1) possesses a N-terminal tail which contains identical residues (RRtVK) [32] and adopts the same binding pattern with the PSI core in the three complexes (Fig. 2c-e). Furthermore, residues from PsaL and PsaH involved in the interactions are also highly conserved during evolution (Fig. 2e). This observation suggests that in *P. patens*, the phosphorylated LHCII itself is able to stably bind to the PSI core, forming the *Pp*PSI-LHCI-LHCII complex similar to the complex found in vascular plants. Formation of the potential *Pp*PSI-LHCI-LHCII complex depends on the phosphorylated LHCII, and does not require Lhcb9[27, 28, 32, 33]. However, compared to the LHCII trimer in *Zm*PSI-LHCI-LHCII complex, LHCII in *Pp*PSI-L complex rotates slightly towards the outer belt and Lhcb9 (Fig. 2c), resulting in additional inter-subunit interactions. These interactions then further stabilize the *Pp*PSI-L complex, which may explain a previous report showing that *Pp*PSI-L is more stable than the *Pp*PSI-LHCI-LHCII complex [32].

### The outer LHCI belt is identical to the inner belt

Our *Pp*PSI-L structure showed that four Lhca proteins form a crescent-shaped outer belt, similar to the inner belt. Interestingly, we found that the second position of the outer belt is also occupied by Lhca2.1 (Fig. S6). In addition, the four outer-Lhcas exhibit the same composition and are arranged in the same order as those in the inner belt. Superposition of the two belts resulted in an RMSD of 1.18 Å, indicating that the relative position of the Lhca proteins is well conserved within the LHCI belts. However, the outer belt of *Pp*PSI-L is slightly less curved compared to the inner belt (Fig. S7a), which may facilitate its association with other parts of the complex (Fig. 3).

**Fig. 3.**
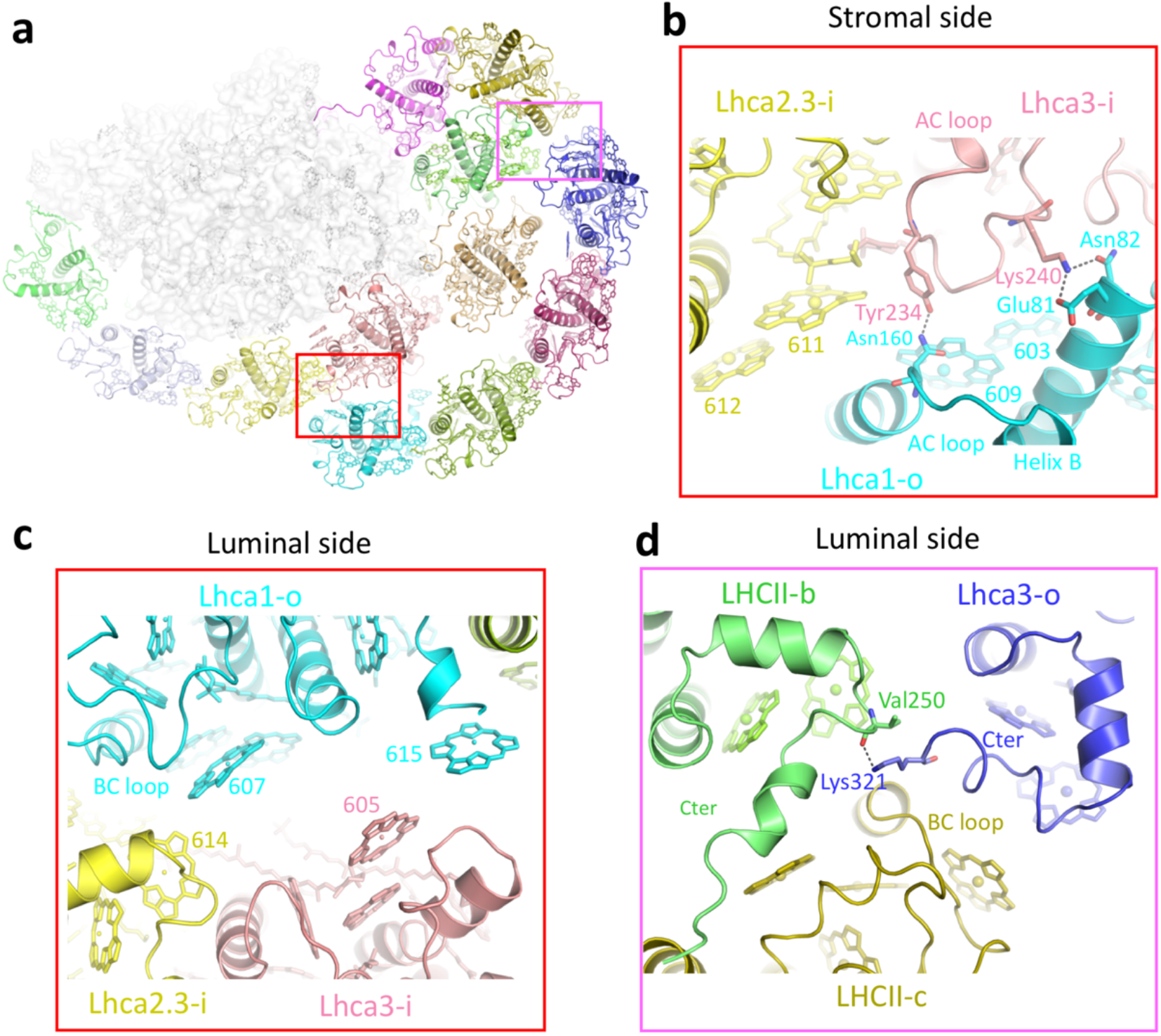
Interactions of the outer LHCI belts with the inner LHCI belt and LHCII. **a**, Cartoon representation of the *Pp*PSI-L. The LHC proteins are shown in the same colors as in Fig. 1a, while PSI core subunits are shown in white color. Interaction regions between the two LHCI belts and between the outer LHCI belt and LHCII are highlighted in red and purple boxes. **b, c**, Detailed interactions of Lhca1-o with Lhca2.3-i and Lhca3-i at the stromal (**b**) and luminal (**c**) side, respectively. **d**, Detailed interactions between Lhca3-o and LHCII. Residues and pigments involved in inter-subunit interactions are labelled and shown in stick mode, the central-Mg atoms of chlorophylls are shown as spheres. Hydrogen-bond interactions are indicated with dashed lines.

Compared to *Cr*PSI-LHCI, the outer belt of *Pp*PSI-L shifts a large distance towards the LHCII trimer, therefore not running parallel with the inner belt. The outer belt interacts with the inner belt and LHCII trimer through the two terminal Lhca proteins (Fig. 3a). The outer Lhca1 (Lhca1-o) interacts with the inner Lhca3 (Lhca3-i) at the stromal side, namely through the N-terminal region of helix B and the AC loop (Fig. 3b). At the luminal side, Lhca1-o forms hydrophobic interactions with the Lhca2.3-i via the BC loop (Fig. 3c). In addition, several Chls (603-609 pair, 607 and 615) of the Lhca1-o form close pairs with pigment molecules of the Lhca2.3-i and Lhca3-i (Fig. 3b, c), further strengthening the association between the two belts. On the other end of the outer belt, Lhca3-o interacts with LHCII primarily at the luminal side, namely through the C-terminal regions of Lhca3-o and LHCII-b, and the BC loop of LHCII-c (Fig. 3d).

When we superposed the outer Lhca1-a2.1-a2.3-a3 belt of the *Pp*PSI-L structure onto the same position of the outer belt (Lhca1-a4-a6-a5) in the *Cr*PSI-LHCI structure, we found that the Lhca1-o and Lhca3-o clash with the inner Lhcas (Fig. S7b). In addition, *Cr*Lhca5 and *Cr*Lhca6 located in the outer belt contain a long C-terminal region, which contributes to the association between the two belts in *Cr*PSI-LHCI (Fig. S7c). In contrast, *P. patens* does not contain homologs of *Cr*Lhca5 and *Cr*Lhca6. Together, these features may result in the shift of the outer LHCI belt in *Pp*PSI-L. Our findings suggest that evolutionary mutations in LHCIs prevent *P. paten*s to form a parallel bilayer LHCI belt similar to that of *C. reinhardtii*, resulting in the specific organization of *Pp*PSI-L.

### The unique protein Lhcb9 plays a central role in the *Pp*PSI–L formation

Our MS result of the *Pp*PSI-L sample identified the presence of Lhcb9.1 in the complex (Fig. S1), which is consistent with previous data showing that Lhcb9.2 is the minor isoform in *P. paten*s [27, 28, 33]. Based on these analyses, we modeled Lhcb9.1 in our structure. Lhcb9 in our model exhibits structural features similar to that of LHCII, namely consisting of three transmembrane helices and two short helices at the luminal side (Fig. 4a). Compared to other LHC proteins, Lhcb9 features a longer N-terminal region and a shorter C-terminal tail (Fig. S8a). Lhcb9 binds 13 chlorophylls (Chl 601-613), but lacks the Chl 614 normally found in LHCII, because that its ligating residue His present in LHCII has been substituted by Trp in Lhcb9. Moreover, Lhcb9 binds four carotenoids, with three located at the conserved binding sites, namely L1, L2 and N1. The fourth carotenoid is located at the molecular surface of Lhcb9, at a site we henceforth call L3, which is unique for Lhcb9 (Fig. S8a). This newly discovered carotenoid is located close to Chl 603, and arranged in parallel to its porphyrin head, with a distance of approx. 4.4 Å (Fig. 4b). The Chl 603 is ligated by an Asn, instead of the usual His used in other Lhcb proteins, and forms a strongly coupled pair with Chl 609, constituting the ‘red Chl’. The red Chl 603-609 is sandwiched by two carotenoids located at the L2 and L3 site, resulting in a pigment cluster composed of two chlorophylls and two carotenoids (Fig. 4b). The particular arrangement of this pigment cluster implies that it plays a role in quenching excess energy in addition to light harvesting.

**Fig. 4.**
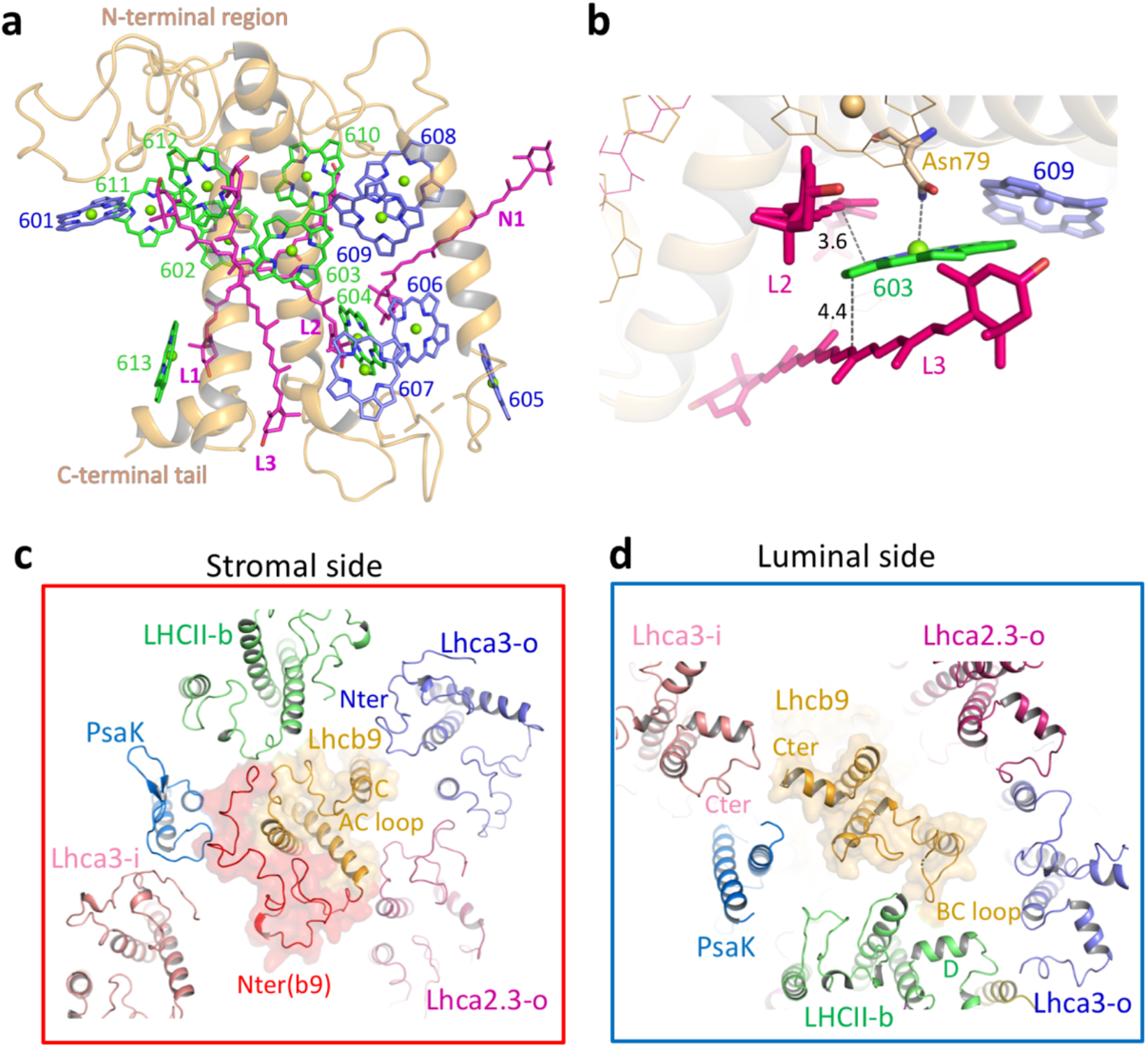
Structure of Lhcb9 and the interactions of Lhcb9 with surrounding subunits. **a**, Cartoon representation of *Pp*Lhcb9. Chls are shown as sticks with the central-Mg atoms shown as spheres. Chls a (green) and Chls b (blue) are assigned according the conserved sites in spinach LHCII (PDB code 1RWT). Carotenoids are shown as sticks and colored purple. **b**, A pigment cluster composed of two chlorophylls and two carotenoids in Lhcb9. Chl 603 is ligated by Asn residue. The edge-to-edge distances between Chl 603 and two carotenoids located at sites L2 and L3 are indicated. **c, d**, Interactions of Lhcb9 with the surrounding subunits viewed from the stromal (c) and luminal (d) side, respectively.

Our structure showed that Lhcb9 is located at the central position of all LHC proteins (Fig. 1a), where it forms close contacts with the PSI core, the LHCII trimer, inner and outer LHCIs. At the stromal side, the long N-terminal tail of Lhcb9 simultaneously interacts with LHCII-b, PsaK and Lhca2.3-o (Fig. 4c). Moreover, its AC loop and helix C form hydrophobic interactions with the AC loop of LHCII-b and the N-terminal region of Lhca3-o. At the luminal side, the BC loop of Lhcb9 associates with helix D of LHCII-b, furthermore, the C-terminal region of both Lhcb9 and Lhca3-i form close contacts (Fig. 4d). The central position of Lhcb9 and the extensive interactions between Lhcb9 and surrounding subunits explained the previous finding that knocking out Lhcb9 causes the completely loss of the *Pp*PSI-L complex[27, 28, 33]. Interestingly, our structure clearly showed that Lhcb9 adopts an opposite orientation compared with other LHCI monomers in the complex. The helix C of Lhcb9 does not orientate towards the Lhca3-i, but is located close to the Lhca3-o and LHCII trimer (Fig. S8b). This orientation of Lhcb9 might be determined by the shapes of both Lhcb9 molecule and the central space where it is located. The long N-terminal fragment of Lhcb9 shapes a convex close to helix A at the stromal side, and is perfectly accommodated by the concave formed by PsaK with neighboring LHCII and Lhca3-i. The specific orientation of Lhcb9 allows its interactions with other LHCs, thus is pivotal for stabilizing the *Pp*PSI-L complex. Moreover, this orientation places the red Chl 603-609 of Lhcb9 facing outward (Fig. S8b), opposite to all Chl 603-609 pairs found in other LHCIs, which are all facing towards the PSI core. Lhcb9 is only found in mosses and absent from both green algae and other plants[23], indicating that Lhcb9 is critically involved in the adaptation of mosses to dim-light terrestrial environments. The specific location and pigment arrangement of Lhcb9 may play an essential part in this function.

### Pigment arrangement and potential EET pathways in *Pp*PSI–L

The *Pp*PSI-L contains a greatly increased number of chlorophyll molecules compared to *Pp*PSI-S, resulting in enhanced light harvesting per PSI unit. Chlorophylls with short distances and belonging to neighboring subunits form multiple interfacial Chl pairs, which contribute to the efficient EET within this complex. Based on our structure, the inner LHCIs, LHCII and Lhcb9 may directly transfer the excitation energy to the core due to their close location to the core subunits. In addition, they may also mediate the EET from outer LHCIs to the core (Fig. 5). The inner LHCI belt has its two terminal Lhca proteins close to the PSI core, whereas Lhca2.1-i and Lhca2.3-i are a little distant from the core. Thus, EET from the inner belt to the core in *P. patens* is probably following similar EET pathways as those in plant PSI-LHCI, i.e. primarily through the terminal Lhca1-i and Lhca3-i [17, 19]. The LHCII trimer binds to the PSI at the similar position as that in *Zm*PSI-LHCI-LHCII complex, hence LHCII may transfer the excitation energy to the PSI core via similar pathways as revealed in *Zm*PSI-LHCI-LHCII structure [38], delivering the excitation energy primarily to PsaO through LHCII-a (LhcbM2) and to PsaK through LHCII-b. Lhcb9 is located adjacent to PsaK, suggesting that Lhcb9 relays the excitation energy directly to PsaK. We found that both Chls 611 and 612 of Lhcb9 are located close to Chl 201 of PsaK, with Mg-to-Mg distances of approx. 12 Å, allowing the highly efficient EET from Lhcb9 to the PSI. In addition, the inner LHCIs, Lhcb9 and LHCII are inter-connected via the interfacial Chl pairs 611_Lhca3-i_-611_Lhcb9_ and 612_LHCII-b_-608_Lhcb9_. This feature suggests that the light energy absorbed by these antenna proteins may quickly equilibrates within the inner LHCI-Lhcb9-LHCII moiety, and is transferred to the PSI through multiple pathways mentioned above.

**Fig. 5.**
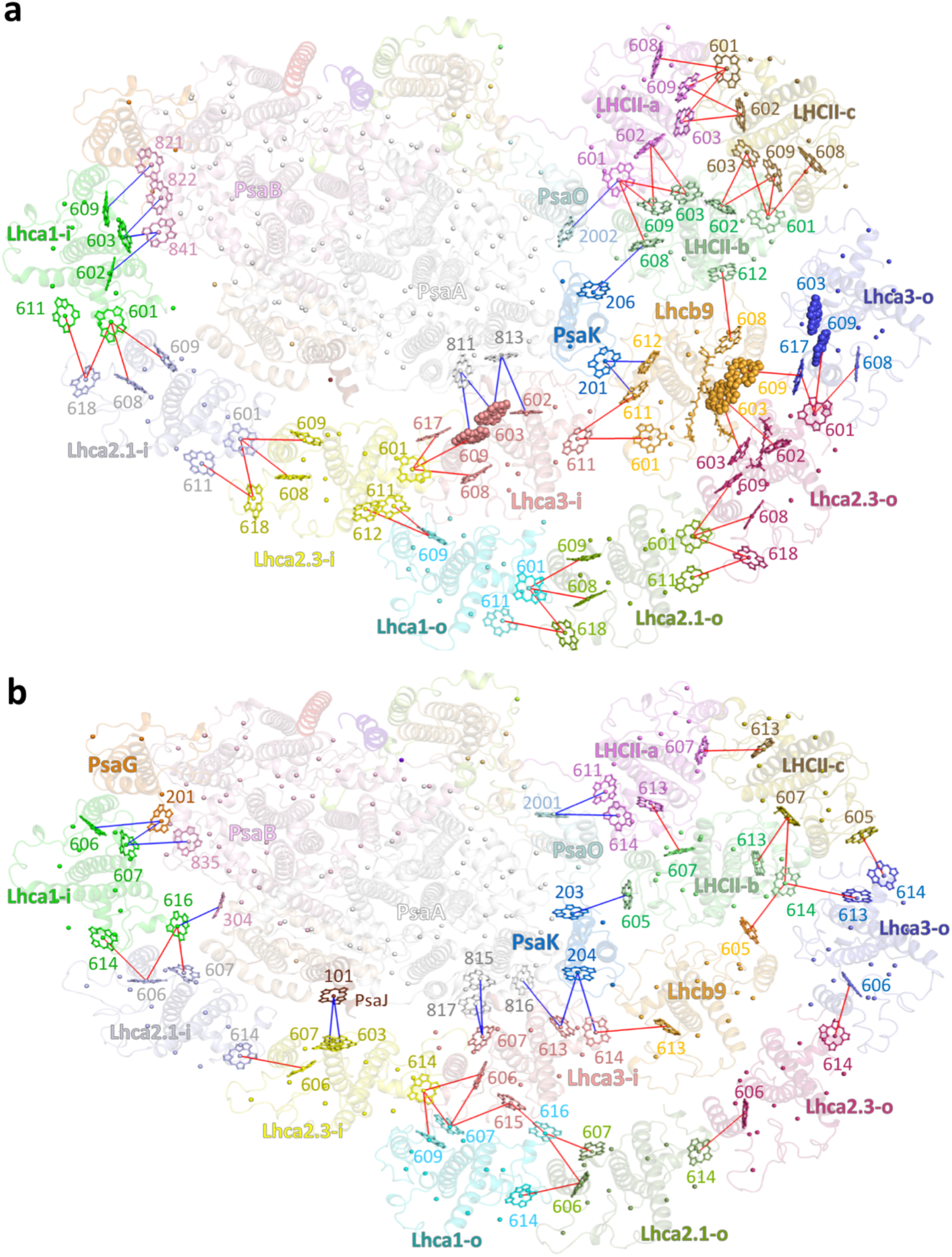
Potential energy transfer pathways in the *Pp*PSI–L complex. **a**, Stromal-side view of chlorophylls within the *Pp*PSI–L complex at the stromal layer (a) and luminal layer (b). Plausible EET pathways between neighboring LHCIs are indicated by red lines, those between LHCIs and the PSI core are indicated by blue lines. Chlorophylls involved in the EET are labeled and shown as spheres for red Chls and sticks for other Chls. For clarity, the phytol chains of chlorophylls are omitted.

The outer LHCI belt binds to the PSI core indirectly thus may transfer the excitation energy to the core through the inner LHCI-Lhcb9-LHCII moiety. Similar to the inner belt, EET from the outer belt towards the core is primarily through the two terminal Lhcas. The Lhca1-o closely interacts with the Lhca2.3-i and Lhca3-i. The EET from Lhca1-o to Lhca2.3-i presumably takes place via two interfacial Chl pairs, the stromal side 609_Lhca1-o_-611_Lhca2.3-i_ and the luminal side 607_Lhca1-o_-614_Lhca2.3-i_, both having a Mg-to-Mg distance of approx. 13 Å. In addition, Lhca1-o may also guide its excitation energy to the Lhca3-i through the luminal Chl pair 616_Lhca1-o_-615_Lhca3-i_. The Lhca3-o is located adjacent to Lhcb9 and LHCII-c. The Chls 617 and 614 of Lhca3-o are close to the stromal side Chl 609 of Lhcb9 and the luminal side Chl 605 of LHCII-c, with Mg-to-Mg distances ranging from 13 to 15 Å. These two Chl pairs may constitute the major EET pathways from Lhca3-o towards the PSI core. The chlorophyll molecules of Lhca2.1-o and Lhca2.3-o are located slightly distant from those of the inner LHCI-Lhcb9-LHCII moiety thus the light energy absorbed by the two outer Lhcas may be firstly transferred to the terminal Lhca1-o and Lhca3-o, and then further towards the core.

### Structural flexibility of *Pp*PSI–L

Large multi-subunit protein complexes usually exhibit structural heterogeneity due to their dynamic conformational variations. These variations play an essential part in ensuring their proper functions. While the *Pp*PSI-L complex is stabilized by multiple inter-subunit interactions, it exhibits internal flexibility as shown by our multi-body refinement and analysis [44]. To perform multi-body refinement, we defined three separate bodies in the *Pp*PSI-L complex, namely the *Pp*PSI-S moiety, the LHCII trimer, and Lhcb9 plus the outer LHCI belt (Lhcb9-outer LHCIs) (Fig. S9a). The result indicated that three major principal components (component 1-3) account for more than 50% of the total variances of the data (Fig. S9b). In the first two components, LHCII, Lhcb9-outer LHCIs appear to move together in a synchronized manner. The first component displays a rotational shift of Lhcb9-outer LHCIs-LHCII moiety relative to *Pp*PSI-S within the membrane plane (Fig. S9c, Movie S1). The Lhcb9-outer LHCIs-LHCII is translated by distances of approx. 10 Å to 13.5 Å at the edges (Lhca1-o and LHCII) and in the middle (Lhca2.3-o). The second component shows a vertical rotation of the Lhcb9-outer LHCIs-LHCII moiety relative to *Pp*PSI-S (Fig. S9d, Movie S2). Lhca2.3-o, which is a distantly located LHC protein from the PSI core, exhibits a profound pivot of 27.4 Å, while Lhca1-o and LHCII at both edges move shorter distances (approx. 9 Å). The third component reveals the separation of LHCII from Lhcb9-outer LHCIs. The two rigid-bodies rotate vertically but towards opposite directions relative to *Pp*PSI-S (Fig. S9e, Movie S3). The mobile behavior of LHCII trimer independently of *Pp*PSI-S and Lhcb9-outer LHCIs suggests that the LHCII trimer alone is able to bind to the PSI core and form *Pp*PSI-LHCI-LHCII complex. Furthermore, our findings indicated that the *Pp*PSI-S, the LHCII trimer and the Lhcb9-outer LHCIs are mobile relative to each other, with the latter two being more closely associated. The dynamic feature of the *Pp*PSI-L complex implies that local rearrangement of the complex may occur under different light conditions. In addition, the relative movement of Lhcb9-outer LHCIs and LHCII in respect to the *Pp*PSI-S alters the interfacial Chl-Chl distances. As a result, the efficiency of EET from LHC proteins to the PSI core is probably changed. Therefore, the dynamic conformational change of *Pp*PSI-L may serve as a regulatory mechanism for fine-tuning the EET within the complex.

### Evolution of Lhca proteins in eukaryotes

During evolution, green algae, mosses and flowering plants emerge in a consecutive manner, transiting from an aquatic towards a terrestrial environment. The PSI antennae of these phototrophs are usually form the LHCI belt composed of four (or sometimes two) Lhcas, attaching to the core complex directly or indirectly. When superposing each Lhca proteins of the (inner) LHCI belt in *P. patens, Z. mays* and *C. reinhardtii* (*Pp*Lhca1-a2.1-a2.3-a3, *Zm*Lhca1-a4-a2-a3 and *Cr*Lhca1-a8-a7-a3) (Fig. S10), we found that while the two terminal Lhcas are highly conserved across evolution, the two middle Lhcas of green algae and land plants show clear conformational variations in their AC loop region. These features are presumably related to the essential roles of the two terminal Lhcas, namely binding with the core and ensuring efficient EET from the LHCI belt to the core [14, 17]. In contrast, the two middle Lhcas are a little distant from the core, playing minor roles in interacting and EET function. Thus, the four Lhcas experienced different levels of evolutionary changes.

In addition, structural comparison showed that the four Lhcas changed their bound chromophores. For example, almost all *Cr*Lhcas (except *Cr*Lhca3 and *Cr*Lhca9) possess a Chl at the position 616[9, 40], which is coordinated by the C-terminus of each Lhca protein. In plants, only Lhca1 preserves this Chl albeit with a small pivot in orientation (Fig. S10a), whereas all other Lhca proteins have lost Chl 616. Chl 616 is located at the Lhca-core interface, where it plays a pivotal role in efficient EET from Lhca to the core in *Cr*PSI-LHCI. Lacking this Chl in land plants may be an acclimate result of experiencing the stronger and fluctuated light conditions in terrestrial environment. In addition, land plants acquired one Chl at the position 618 in the two middle Lhcas. Structures showed that the Chl 618 is coordinated by the AC loop, which adopts different conformation between green algae and land plants. The Chl 618 is located at the interface between two neighboring Lhcas and on the exterior surface of the complex, implying that plants evolve Chl 618 to strengthen the interaction between neighboring Lhcas and enhance their light harvesting capability upon losing the outer LHCI belt. Moreover, Lhca4 of flowering plants binds an additional Chl at the position 617 which is ligated by a His located in helix C. This His residue is not conserved in *P. patens* and *C. reinhardtii*, indicating that this His evolved later, presumably for binding Chl 617. Adjacent to Chl 617, one extra carotenoid is located at the Lhca1-a4 interface. Thus, the newly evolved Chl 617 and the carotenoid increase the association between Lhca1 and Lhca4 and the interaction of Lhca1-Lhca4 with the core in vascular plants. Lhca3 shows the opposite feature to Lhca4, binding decreased number of carotenoids from green algal Lhca3 to plant Lhca3. Green algal Lhca3 contains five carotenoids, located at the three conserved binding sites (L1, L2 and N1), and two other binding sites at the molecular surface, facing the core (Car1) and the outer belt (Car2). *Pp*Lhca3 loses Car1, but preserves Car2, which may be pivotal for the binding of the outer belt in *P. patens* and *C. reinhardtii*. In agreement with this assumption, Lhca3 of flowering plants lack both carotenoids together with having lost the outer LHCI belt.

## Discussion

Here, we report the near-atomic resolution structure of the *Pp*PSI-L complex, and describe the composition and interactions of the protein subunits as well as the sophisticated network of pigments associated with this complex. We demonstrated that Lhca5 is absent in our *Pp*PSI-S and *Pp*PSI-L samples, suggesting that Lhca5 only exist in the NADP(H) dehydrogenase-associated PSI, similar to its counterpart in vascular plants (*A. thaliana* and *Hordeum vulgare*) [45, 46]. In agreement with previous reports [27, 33], we showed that Lhcb9 is located at the central position of LHC proteins, forming multiple interactions with the PSI core, LHCII trimer and LHCIs from both the inner and outer belts, explaining the previous finding that the *Pp*PSI-L complex disassembled in the absence of Lhcb9. We found that *Pp*PSI-L contains inner and outer LHCI belts with identical composition and arrangement of Lhca1-a2.1-a2.3-a3. In contrast to *Cr*PSI-LHCI, the outer LHCI belt in *Pp*PSI-L complex does not run parallel to the inner one, but moves towards the direction of LHCII, with only one LHCI (Lhca1-o) interacting with the inner Lhca proteins. This organization pattern results in an unstable outer LHCI belt, explaining why in flowering plants, this belt is lost as a result of missing Lhcb9. However, in mosses that contain Lhcb9, this assembly allows the accommodation of Lhcb9 in the central space and additional interactions of the outer LHCI with Lhcb9 and the LHCII trimer, most likely further stabilizing the *Pp*PSI-L complex.

Previous reports revealed that Lhcb9 is expressed in low-light conditions and increases the functional antenna size of *Pp*PSI [27, 33]. The specific organization of the Lhcb9-mediated *Pp*PSI-L complex allows all LHC proteins to be linked with each other. This organization results in a highly connected chlorophyll network which may be critical for the efficient EET within the complex. In addition, the *Pp*PSI-L complex contains three Lhcas that carry red chlorophylls, namely Lhcb9 and two Lhca3 subunits. The red Chls are able to harvest long-wavelength light which are enriched under shade conditions, thus may play a critical role for light harvesting in mosses. In *P. patens*, these red Chls are distributed at the central region of the Chl network (Fig. 5), suggesting that they contribute to the efficient EET in addition to increasing the absorption cross-section. Moreover, the red Chl 603-609 of Lhcb9 and Lhca3.1-o are located in close vicinity (Fig. 5), resulting in a red Chl cluster that might be critical for protecting *P. patens* PSI from damage under stress conditions. A recent report showed that the accumulation of guanosine tetraphosphate and pentaphosphate ((p)ppGpp), which are important nucleotides in stress acclimation, was accompanied by an increase in Lhcb9 levels [47], suggesting that the Lhcb9-mediated *Pp*PSI-L complex plays a role in coping with environmental stresses.

In addition, our findings demonstrated that LHCII within the *Pp*PSI-L complex is phosphorylated on at least one monomer, which we assigned as LhcbM2. LhcbM2 interacts with the *Pp*PSI-S part in a pattern identical to that of *Zm*PSI-LHCI-LHCII. Moreover, the *Pp*LHCII trimer pivots towards the stroma in relation to the PSI core, a feature also observed in the *Zm*PSI-LHCI-LHCII structure [38]. These observations, together with our multi-body analysis, strongly suggests that the *Pp*PSI-LHCI-LHCII complex exhibits the same architecture as the plant counterpart. Based on our results, we proposed that the three types of *Pp*PSI (*Pp*PSI-S, *Pp*PSI-LHCI-LHCII and *Pp*PSI-L) are formed by the same building blocks, namely the PSI core, the Lhca1-a2.1-a2.3-a3 belt, and optionally the phosphorylated LHCII trimer and Lhcb9. During the PSI assembly process, four Lhcas are presumably forming the LHCI belt first, which then associates with the PSI core [48]. Thus, a portion of free LHCI belts may exist and could either bind to the core or serve as the outer belt in the presence of Lhcb9. *P. patens* represents an evolutionary intermediate between green algae and flowering plants, and grows at the interface between aquatic and terrestrial environments. The highly fluctuating light conditions that *P. patens* faces require the fast and dynamic adjustment of the *Pp*PSI antenna size. This requirement may be fulfilled by using these shared modules in multiple types of *Pp*PSI. The *Pp*PSI-S might be the major PSI form in *P. patens* in normal light. Under state 2 conditions, LHCII is phosphorylated and binds to the *Pp*PSI-S, forming the *Pp*PSI-LHCI-LHCII complex. Upon prolonged low light acclimation, *P. patens* increases the expression level of Lhcb9, which further recruits the LHCI belt to associate with *Pp*PSI-LHCI-LHCII, resulting the formation of *Pp*PSI-L. The curved arrangement of the *Pp*PSI-L (Fig. 1) may facilitate the quick switch between *Pp*PSI-S, *Pp*PSI-LHCI-LHCII and *Pp*PSI-L upon a slight alteration of the thylakoid membrane architecture. Thus, the three types of PSI achieve a dynamic balance in *P. patens*, and adjusting their ratios depending on different conditions likely represents a critical regulatory mechanism for light harvesting in *P. patens*.

## Methods

### *Pp*PSI-L purification

Moss *P. patens* was grown on plates for 14 days under continuous white light (40-50 µmol photos m^-2^s^-1^ for the first ten days, and 17 µmol photos m^-2^s^-1^ for the last four days) using standard BCDAT media in the presence of 0.5% glucose [28, 33]. In addition, *P. patens* grown under the same conditions without glucose were used as control. Plants were scraped off the plates, and the protonemal tissues were ground using Waring blender in a buffer containing 20 mM Tricine-KOH (pH 7.8), 400 mM NaCl, 2 mM MgCl_2_, 1 mM PMSF, 0.2 mM benzamidine, and 1 mM *ε*-aminocaproic acid at 4 °C. The suspension was filtered through 4 layers of cheesecloth and centrifuged at 27,000 х g for 10 min. The pellet was resuspended in a buffer containing 20 mM Tricine-KOH (pH 7.8), 150 mM NaCl, 5 mM MgCl_2_, 0.2 mM benzamidine, and 1 mM *ε*-aminocaproic acid [28]. The thylakoid membrane fractions were further purified by sucrose cushion ultracentrifugation method [24]. In short, thylakoids were resuspended in 1.8 M sucrose, 5 mM Tricine (pH 7.5), 10 mM EDTA overlaid by solution containing 1.3 M / 0.5 M sucrose, 5 mM Tricine (pH 7.5), 10 mM EDTA, and centrifuged at 103,900 х g for 1 h at 4 ℃ using a P28S rotor (HATACHI). After centrifugation, the purified membranes concentrated at the interface of 1.3 M / 1.8 M sucrose cushion were extracted, and diluted with buffer containing 5 mM Tricine Ph 7.5, and then centrifuged at 27,000 х g for 10 min. The pellet was resuspended in a small volume of buffer containing 20 mM Tricine (pH 7.5), 10 mM NaCl, 5 mM MgCl_2_ and 400 mM sucrose and stored at -80 ℃for future use.

For purification of the *Pp*PSI-L complex, the thylakoid membranes were diluted to a concentration of 0.5 mg chlorophyll mL^-1^ and solubilized with 1% (w/v) dodecyl-β-D-maltoside (β-DDM) (Anatrace) for 30 min on ice. The solubilized membranes were then loaded onto the 0.1-1.3 M sucrose density gradient in a buffer containing 25 mM MES-NaOH (pH 6.5) and 0.03% (w/v) dodecyl-α-D-maltoside (α-DDM), following centrifugation at 154,390 × g for 20 h using SW41 rotor (Beckman) at 4 °C [28]. The green bands corresponding to *Pp*PSI-L and *Pp*PSI-S were harvested.

### Characterization of the *Pp*PSI-L

Absorption spectra of *Pp*PSI-L and *Pp*PSI-S samples were recorded using spectrometer (U-3900, HATCHI, Japan) (Fig. S1c). Pigment analysis (Fig. S1d) was performed by high-performance liquid chromatography (HPLC) using Hitachi Elite LaChrom HPLC System. The *Pp*PSI-L and *Pp*PSI-S samples after sucrose density gradient centrifugation were separately mixed with 80% (v/v) cold acetone, and centrifuged at 13,000 × g for 15 min to extract pigments. The supernatant was loaded on an Allsphere ODS-2 column and the elution was detected by Hitachi L-2450 Diode Araay systems (HATACHI, Japan). The elution protocol was the same as that described previously [49]. Pigments were identified based on their absorption spectra and elution times. The *Pp*PSI-L and *Pp*PSI-S samples were analyzed by SDS-PAGE (Fig. S1b) [50]. Each single band in the SDS-PAGE was cut off for identification of the protein subunit through mass spectrometry (MS) analysis. MS analysis (Source data) was performed using a matrix-assisted laser desorption ionization time-of-flight mass spectrometer MALDI-TOF (UltrafleXtreme, Brucker, Germany).

Immunoblot analysis (Fig. S1e) was performed according to the previous report [51]. *Pp*PSI-L, *Pp*PSI-S and thylakoid membrane samples were fractionated by SDS-PAGE and electroblotted onto Polyvinylidene-Fluoride (PVDF) membranes. Lhcb9 and phosphorylated LHCII were detected using antibodies of α-Lhcb9 (AS15 3088) from Agrisera and α-Thr-P (9381S) from Cell Signaling Technology (CST), respectively. PsaA was used as a control and detected by α-PsaA (AS06 172) from Agrisera.

### Cryo-EM sample preparation and data acquisition

The *Pp*PSI-L sample after sucrose gradient density was buffer exchanged with 25 mM MES pH 6.5, 10 mM NaCl, 5 mM MgCl_2_ and 0.5 M betaine, and concentrated to 2 mg chlorophyll mL^-1^ using a 100-kDa molecular-weight cut-off centrifugal filter unit (Amicon Ultra-15, Merck Millipore). A 3 µl volume of the sample was applied to a glow-discharged holey carbon grid (GIG, Au R1.2/1.3, 300 mesh), with a blotting time of 3 s, blotting force of level 3, 100% humidity and 4 °C, and plunge-frozen in liquid ethane using a Vitrobot device (Thermo Fisher Scientific).

All micrographs of *Pp*PSI-L were collected using SerialEM software [52] on a 300 kV Titan Krios electron microscope (Thermo Fisher Scientific) equipped with a Gatan K2 Summit direct detector camera and a GIF quantum energy filter (20 eV). A total of 7455 images from three independent data sets were collected, each was exposed for 6 s and dose fractionated into 32 frames, giving a total electron dose of ∼ 60 e^-^Å^-2^. Data were collected in a defocus range of 1.5-2.5 µm at ×130,000 magnification and a pixel size of 0.52 Å in super-resolution mode.

### Data processing

All movies collected were aligned, dose weighted and summed using MotionCor2 [53] and binned to a pixel size of 1.04 Å. The contrast transfer function (CTF) parameters for the summed images were determined by the program CTFFIND4 [54]. The data processing steps were performed using the software RELION v.3.0 [55] including reference-based auto-picking, 2D/3D classification, 3D auto-refine, CTF refinement and Bayesian polishing. 291,778, 487,775 and 207,857 particles were picked up from the three data sets. After the 2D classification, 163,874, 315,726 and 85,789 particles were kept and then combined for subsequent 3D classification and refinement. To further improve the density of the outer LHCIs, Lhcb9 and LHCII, focused refinement was performed with a soft-edged mask around the PSI-LHCI, outer LHCIs plus Lhcb9, LHCII plus PsaH, PsaO and PsaL regions. Local maps were generated in UCSF Chimera [56], and were further processed into masks using RELION [55]. Afterwards, the overall and the three local density maps were optimized separately by Bayesian polishing, resulting in the final resolutions of 2.87 Å, 2.82 Å, 3.29 Å and 3.05 Å respectively, estimated based on the gold-standard Fourier shell correlation with the 0.143 criterion. Moreover, the density of the outer LHCIs plus Lhcb9 was further optimized using DeepEMhancer software [57]. The specific data-processing workflow is shown in the Fig. S2. The local resolution of the final map was calculated using ResMap [58].

### Model building and refinement

To construct a structural model of the *Pp*PSI-L complex, the structure of *Pp*PSI-S complex (PDB: 7KSQ) was docked into the 2.87 Å resolution map and the 2.82 Å focused refinement map using UCSF Chimera [56]. The LHCII moiety was built using the Lhcb2 of *Zm*PSI-LHCI-LHCII (PDB: 5ZJI) as initial model and docked into a 3.05 Å focused refinement map. The unique protein, Lhcb9, was built using LhcbM5 of *Cr*PSI-LHCI-LHCII (PDB: 7D0J) as initial model, and outer LHCIs were modeled with LHCIs in *Pp*PSI-S structure (PDB: 7KSQ), both of them were docked into a 3.29 Å focused refinement map. Each subunit was mutated with Chainsaw in CCP4 [59] based on the amino acid sequence of the *P. patens* genome. The overall model was rebuilt and adjusted manually using Coot [60], and combined the three focused refinement maps into a composite map for overall refinement with Phenix v.1.19.2 [61], which provided geometric constraints on the cofactors and Chl-ligand relationships. Automatic real-space refinement using Phenix and manual correction using Coot were performed interactively. Statistics on data collection and structural optimization of *Pp*PSI-L complex are summarized in Table S1. The geometries of the final structures were assessed using MolProbity [62]. High resolution images for publication were produced using Chimera and PyMOL (Molecular Graphics System).

### Multi-body refinement and analysis

To evaluate the mobility of LHCII trimer and outer LHCIs plus Lhcb9 relative to the *Pp*PSI-S moiety in the *Pp*PSI-L complex, the refined particles obtained after CTF refinement and Bayesian polishing procedures were subjected to multi-body refinement [44, 63] in RELION v.3.1. Three rigid bodies, corresponding to *Pp*PSI-S moiety, the LHCII trimer and Lhcb9-outer LHCIs were assigned by individual masks. Initial angular sampling rate, offset range and offset step were set to 7.5°, 6 pixels and 1 pixel, respectively. The converged refinement yielded an overall resolution of the map at 3.36 Å. In the subsequent flexibility analysis through the relion_flex_analyse programme, the motion of the three rigid bodies was decomposed along 18 different eigenvectors. The principal components along the top three eigenvectors were output as movies, each comprising ten maps (that is, ten frames for each video). Bin 1 represents the first frame of the movie while bin 10 represents the last. These two frames correspond to the two extreme positions of motion along a specific eigenvector. The top three eigenvectors account for 26.3%, 14.9% and 11.4% of variance in the rotations and translations of the three bodies, respectively. The structural models of *Pp*PSI-S moiety, the LHCII trimer and Lhcb9-outer LHCIs were fit into the bin 1 and bin 10 maps of the specific component in the Chimera program and verified in Coot.

### Identification of Lhca2 and LhcbM proteins

In the structure of *Pp*PSI-L, the two subunits were assigned as Lhca2.1 and Lhca2.3 in both inner and outer LHCI belts and the LHCII-a monomer was assigned as LhcbM2, according to the specific density features of masked maps. We were unable to identify LHCII-b and LHCII-c monomers and thus the two subunits were tentatively built using LhcbM2 sequence. For the identification of Lhca2.1 in both inner and outer LHCI belts, the map features of M103 and F237 exclude the possibility of other Lhca2 isoforms and Lhca5 protein.

The densities of other characteristic residues, including L69, W77, W113 and W163, were used to further verify the assignment of both Lhca2.1-i and Lhca2.1-o (Figs. S3a, S6a). For the identification of Lhca2.3 in the inner LHCI belts, the map features of A150 and R222 exclude the possibility of other Lhca2 isoforms and Lhca5 protein. The densities of other characteristic residues, including F69, F103, V238 and Y243, were used to further verify the assignment of Lhca2.3-i (Fig. S3b). In the outer LHCI belt, the map features of Q73, P74 and D218 exclude the possibility of other Lhca2 isoforms and Lhca5 protein. The densities of other characteristic residues, including residues F69, F103, V238 and Y243, were used to further verify the assignment of Lhca2.3-o (Fig. S6b).

For the identification of LhcbM2 in the LHCII trimer, the LHCII-a is clearly phosphorylated at its N-terminal tail by analyzing the map feature of residues R38, R39 and pT40 (Fig. S5c). Moreover, we were able to narrow down the candidates for the phosphorylated monomer to LhcbM1, LhcbM2, LhcbM4, LhcbM5 and LhcbM13 by analyzing the feature of residues Y57 instead of F57 of our cryo-EM map (Fig. S5d). Furthermore, our MS analysis of the *Pp*PSI-L sample detected unique peptides belonging to LhcbM1 and LhcbM2. In addition, we found that the N-terminal tail of LHCII-a adopts the same conformation and binds at the similar position as that in *Zm*PSI-LHCI-LHCII (PDB: 5ZJI) and *Cr*PSI-LHCI-LHCII (PDB: 7D0J), and the phosphorylated LHCII monomer in the three structures has the first five residues identical to each other (RRtVK). Based on these results, we assigned the LHCII-a monomer to be LhcbM2.

## Data availability

The atomic coordinate of the *Pp*PSI-L complex has been deposited in the Protein Data Bank with an accession code of 8HTU. The cryo-EM map of the complex has been deposited in the Electron Microscopy Data Bank with an accession code of EMDB-35018. Source data are provided with this paper. All other data generated or analyzed are available from the corresponding authors on reasonable request.

## Acknowledgements

We thank Y. K. He and C. L. Ju from College of Life Science, Capital Normal University, for providing the *P. patents* strain; L. H. Chen, X. J. Huang, B. L. Zhu and F. Sun at the Center for Biological Imaging (IBP, CAS) for the support in cryo-EM data collection; The project is funded by the Strategic Priority Research Program of CAS (XDB27020106, XDB37020101), National Natural Science Foundation of China (31930064 and 31970264), and supported by the National Laboratory of Biomacromolecules (2020kf05).

## Author contributions

X.P. and M.L. conceived and coordinated the project. H.Sun and H.Shang performed the purification and characterization of the *Pp*PSI-L sample; H.Sun and X.P. processed the cryo-EM data, built and refined the structural model. H.Sun performed the multi-body refinement. H.Sun, X.P. and M.L. analyzed the data and wrote the manuscript; all authors discussed and commented on the results and the manuscript.

